# vartracker: an end-to-end tool for pathogen longitudinal variant analysis and visualisation

**DOI:** 10.64898/2026.05.06.723370

**Authors:** Charles S.P. Foster, William D. Rawlinson

**Affiliations:** Serology and Virology Division (SAViD), NSW Health Pathology, Prince of Wales Hospital, Sydney, NSW 2031, Australia; School of Biomedical Sciences, Faculty of Medicine & Health, University of New South Wales, Sydney, NSW 2052, Australia; School of Clinical Medicine, Faculty of Medicine and Health, University of New South Wales, Sydney, NSW, Australia

**Keywords:** pathogen, longitudinal sequencing, Python, open source

## Abstract

Longitudinal sequencing can reveal fine-grained pathogen evolution during acute and chronic infections and inform public health responses. However, integrating ordered pathogen genomic data into a coherent evolutionary and clinical framework can be tedious and error-prone. We present vartracker, an open-source tool for longitudinal pathogen variant analysis and visualisation. Given an ordered sample manifest, vartracker supports three entry points: raw sequence reads, reference-aligned BAM files, or user-supplied VCF and coverage inputs. Raw-read and BAM inputs are processed through an integrated Snakemake workflow, whereas VCF mode starts from precomputed files. Variants are normalised and annotated relative to a reference genome, tracked across timepoints, and classified as original or newly emerging and as transient or persistent. Inferred amino acid changes are reported, and for SARS-CoV-2 analyses, relevant published literature for key mutations can be automatically linked through a functional database. vartracker outputs a schema-documented results table, provenance metadata for reproducibility, publication-quality static figures, and an interactive heatmap for data exploration. Although packaged with SARS-CoV-2 reference assets and initially developed for SARS-CoV-2 datasets, vartracker is pathogen-agnostic when appropriate reference data are supplied. We demonstrate its utility using SARS-CoV-2 and respiratory syncytial virus A (RSV-A) datasets. vartracker is freely available through GitHub, PyPI and Bioconda.

## Introduction

Whole-genome sequencing (WGS) is a vital component of global public health surveillance. Routine WGS at a population level enables pathogen evolution to be inferred and tracked over time, which allows the inference of transmission networks and mutations that might drive the differential success of pathogen lineages [1]. At smaller scales, longitudinal WGS of pathogens from chronically infected individuals and/or *in vitro* serially passaged samples can uncover within-host evolutionary dynamics [2,3]. Linking variants with disease phenotype and clinical outcome metadata can support hypotheses about how mutations influence patient prognosis and disease progression.

There is an abundance of pipelines that tie together the use of bioinformatics tools into easily executable end-to-end workflows for pathogen WGS (Table 1). Population-surveillance platforms such as CoV-Spectrum [4], VIRUS-MVP [5] and the Nextstrain ecosystem [6] operate principally on consensus sequences aggregated across populations, whereas workflows such as nf-core/viralrecon [7] and V-pipe [8] focus primarily on upstream read processing and variant calling. However, interpretation and visualisation of variant calls across serial samples remains a practical bottleneck in longitudinal pathogen sequencing studies because of a paucity of available published tools dedicated to this purpose. Valuable earlier examples include FLEA [9], LAVA [10] and VIPERA [11], but these tools are each relatively specialised: FLEA is tailored to long-read full-length HIV env amplicons, LAVA remains available primarily as a 2019 preprint with a workflow-oriented implementation and no formal GitHub releases, and VIPERA is designed specifically for analysis of FASTA or BAM files from intra-host serially sampled SARS-CoV-2 infection. In practice, longitudinal WGS studies will instead use in-house scripts or *ad hoc* workflows per study to explore the trends in the data (e.g. [12]), which hinders reproducibility.

**Table 1.** A comparison of a selection of tools and workflows for genomic surveillance and longitudinal analysis of virus whole-genome sequencing data.

| Tool | Primary focus | Accepted inputs | Variant calling | Longitudinal / minor-variant analysis | Annotation / context | Visualisation | Repository status* |
| --- | --- | --- | --- | --- | --- | --- | --- |
| **Vartracker**<br>(this study) | Pathogen-agnostic comparison of variants across ordered samples | Illumina FASTQ or BAM; pre-called VCF + coverage (any platform) | **Integrated:** LoFreq for Illumina; other platforms require an external caller and VCF mode | **Direct:** emergence, persistence, fluctuation and loss; retains AF, depth and quality | AA consequences; optional literature-aware SARS-CoV-2 annotation | AF trajectories; heatmaps; position plots; turnover; lifespan; cumulative and gene summaries | **Active:** 3 Aug 2026; v2.3.0 (3 Aug 2026) |
| **FLEA** [9] | HIV env haplotypes, phylogeny and selection | PacBio full-length env amplicons | **Specialised:** error correction and haplotype reconstruction | **Direct:** timepoint haplotype and motif dynamics; HIV env only | Selection; motifs; phylogeny; protein-structure context | Alignment, haplotype-frequency, motif, MDS and structure views | **Inactive:** last updates 2017-2018; no formal release |
| **LAVA** [10] | Longitudinal viral deep-sequencing workflow | Trimmed FASTQ; reference; time metadata | **Integrated:** BWA-MEM and VarScan | **Direct:** allele-frequency trajectories over time | ANNOVAR | HTML report with AF, coverage and trajectory plots | **Inactive:** 18 Aug 2023; no formal release |
| **VIPERA** [11] | Serial or chronic SARS-CoV-2 infection | Sorted BAM; consensus FASTA; metadata | **External:** requires paired preprocessed inputs | **Direct:** rich within-host analysis; SARS-CoV-2 only | Lineage admixture; phylogeny; diversity; selection | Accumulation plots; AF trajectories; heatmaps; trees; diversity | **Active:** 22 Jul 2026; v1.3.1 (2 Jan 2026) |
| **nf-core/viralrecon** [7] | Assembly, consensus and low-frequency calls | Raw reads | **Integrated:** configurable aligners and callers | **Partial:** per-sample calls; no ordered-series model | Pangolin, Nextclade and pathogen-specific outputs | MultiQC and per-sample reports; no serial AF plots | **Active:** 27 Jul 2026; v3.0.0 (28 Oct 2025) |
| **V-pipe** [8] | Within-sample SNVs, diversity and haplotypes | Short- or long-read data | **Integrated:** mapping, calling and haplotype reconstruction | **Partial:** within-host diversity; no ordered-series summary | **Limited:** variant and haplotype outputs | Primarily analysis and QC; longitudinal plots not central | **Active:** 15 Jul 2026; v3.0.0.pre1 (10 Jun 2024) |
| **HaROLD** [13] | Longitudinal whole-genome haplotype reconstruction | Mapped reads; variant-frequency data | **External:** consumes existing variant frequencies | **Direct:** cross-sample covariation; reconstruction-focused | **Limited:** no integrated biological or literature annotation | **None:** no comprehensive integrated suite | **Inactive:** 13 Feb 2023; v2.0.0 (28 Mar 2022) |
| **CoV-Spectrum and LAPIS API** [4,14] | Population-level SARS-CoV-2 surveillance and API | Public consensus sequences; metadata | **None:** no read-level calling | **Population:** trends over time and geography, not within-host | Lineages; mutations; geographic and epidemiological context | Prevalence, time, geography and mutation plots | **Active:** 16/27 Jul 2026; LAPIS v0.8.3 (28 Apr 2026) |
| **VIRUS-MVP** [5] | Population mutation explorer | Processed genomic and metadata outputs | **External:** companion nf-nCoV-voc workflow | **Population:** prevalence trends, not within-host trajectories | Lineage; place; time; curated function | Mutation heatmaps and stratified prevalence | **Active:** 21 Jul 2026; v1.0.0-no-upload (28 Aug 2025) |
| **Nextstrain ecosystem (including Nextclade)** [6,15] | Consensus QC, clades, phylogeny and outbreaks | Consensus FASTA; metadata | **None:** no read-level calling | **Population:** evolutionary time, not within-host AF | Clades; mutations; QC; phylogenetic and epidemiological context | Trees; maps; frequencies; entropy; mutation views | **Active:** 24 Jul 2026; v3.21.2 (28 Apr 2026) |

Direct = explicitly analyses serial or ordered samples; Partial = relevant within-host outputs without a dedicated ordered-series model; Population = temporal comparisons between populations rather than within-host trajectories. \*Repository activity checked 3 Aug 2026; Active denotes a default-branch update in the preceding 12 months. AF, allele frequency; AA, amino acid; MDS, multidimensional scaling.

To fill this analytical gap, we developed vartracker, a lightweight and flexible tool for longitudinal analysis, annotation and visualisation of pathogen variation across serial samples. Although initially developed to investigate the evolutionary trajectories of SARS-CoV-2 isolates during long-term serial passaging [3], vartracker can also support longitudinal analyses in individual clinical cases and public health investigations, and is pathogen-agnostic when suitable reference data are provided. The tool was designed to simplify tracking of variants across serial samples by summarising and annotating patterns such as variant gain and loss, persistence, and changes in allele frequency. It produces rich outputs describing variant trajectories across samples, customisable plots of temporal trends, and optional links to relevant published literature when user-supplied reference annotations are available. vartracker supports multiple entry modes: an end-to-end workflow that processes raw short reads through an integrated Snakemake pipeline, a mode that accepts per-sample alignments against a chosen reference genome (BAM format), and a mode that accepts precomputed per-sample variant call files (VCF) together with reference-wide depth of coverage information. To support reproducible analysis, vartracker generates schema-documented tabular outputs and run-level provenance metadata, and is available as versioned releases via GitHub, PyPI, Biocontainers and Bioconda.

In this manuscript, we describe the implementation, usage and representative applications of vartracker using SARS-CoV-2 and respiratory syncytial virus A (RSV-A) as examples. By transforming ordered sequencing data into standardised, interpretable summaries of variant emergence, persistence, turnover and allele-frequency change through time, vartracker enables reproducible longitudinal analysis and exploration of pathogen evolution while reducing reliance on bespoke downstream scripting.

## Materials and Methods

### Overview and implementation

vartracker was designed with the goal of simplifying the longitudinal tracking and summarisation of variants (SNPs and indels), including inferring the functional impact of variants via links to published literature (Figure 1). The tool can be deployed via three main entry points: (1) ‘vartracker vcf’, when users have (per sample) a variant-call format (VCF) file with variants against a reference genome and a file with per-site depths; (2) ‘vartracker bam’, when users have (per sample) a BAM-format file with alignments of sequencing reads against a reference genome; and, most generally, (3) ‘vartracker end-to-end’, when users want reproducible end-to-end analysis of raw sequencing reads (Figure S1). The core vartracker framework incorporating variant tracking and annotation stages (see below) operates on normalised variant and coverage inputs and is therefore largely sequencing-platform agnostic.

**Figure 1.**
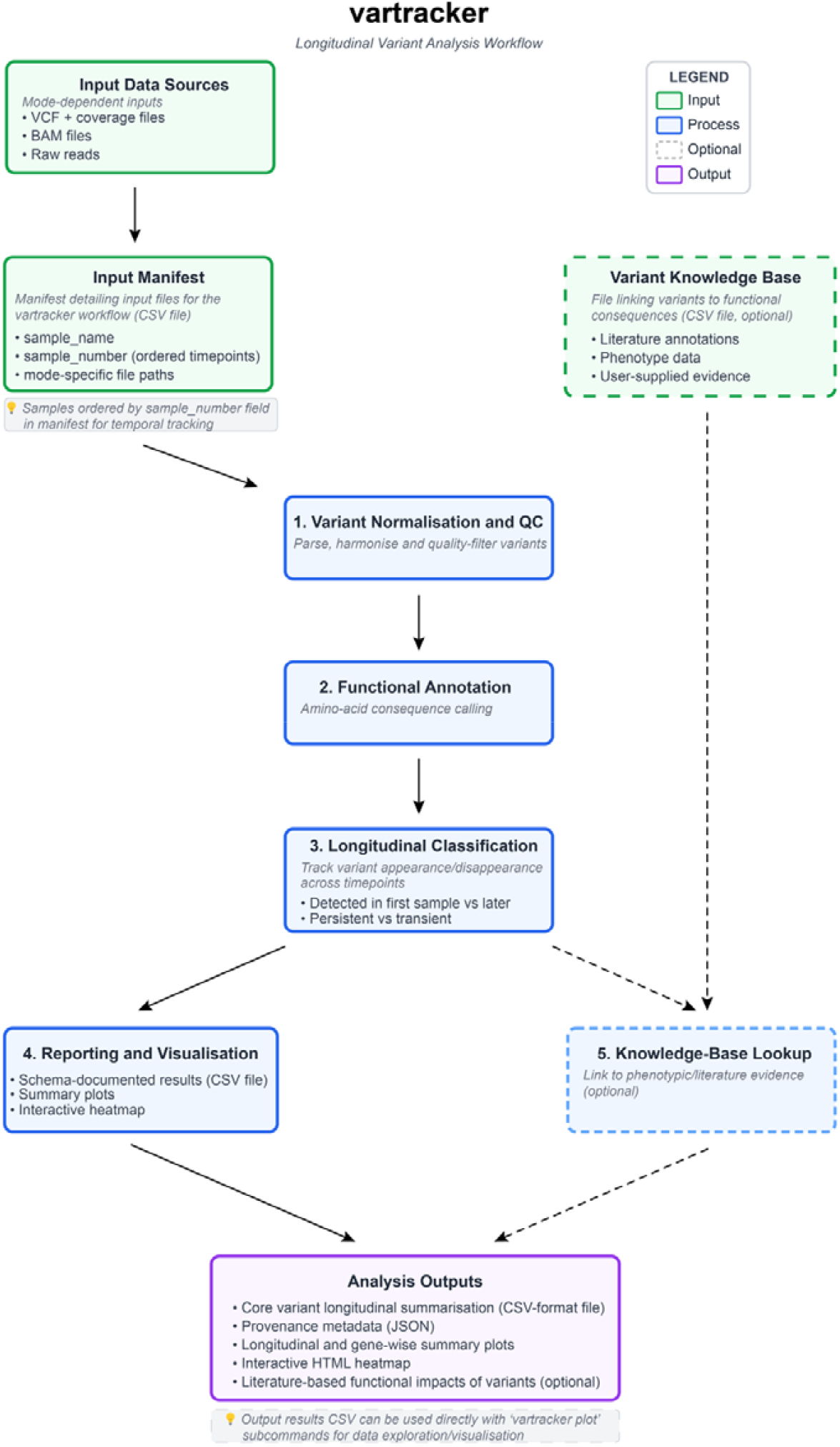
Overview of the vartracker workflow. Conceptual overview of the vartracker longitudinal variant analysis workflow. vartracker accepts multiple input modes, including VCF and coverage files, BAM files, or raw sequencing reads, together with a manifest defining sample order across serial timepoints. Variants are normalised and quality controlled, functionally annotated, and classified according to their longitudinal behaviour across samples. Optional knowledge-base lookup links variants to user-supplied or literature-derived functional evidence. Outputs include schema-documented result tables, provenance metadata, summary plots, and an interactive heatmap.

Vartracker applies configurable minimum allele-frequency and read-depth thresholds when summarising and visualising longitudinal variant calls. The default values were selected based on their use in our genomic surveillance and longitudinal sequencing workflows and should be regarded as starting points rather than universally applicable QC recommendations. Appropriate thresholds depend on the study objective and on the sequencing protocol, depth, variant caller and empirically established error profile of the upstream workflow. More stringent thresholds should be considered when specificity is prioritised or the input data have higher error rates, whereas lower thresholds should be used for low-frequency variant analysis only when supported by a suitably validated upstream workflow. Additionally, the optional integrated end-to-end workflow and BAM-based processing were developed and evaluated primarily for Illumina short read data. Results derived from long read data should be processed using an appropriate platform-specific variant caller before analysis with the ‘vartracker vcf’ input mode, provided that appropriate upstream mapping, variant calling and filtering have been performed.

With the exception of some components of the end-to-end workflow, vartracker is written in Python. Depending on the entry mode, vartracker calls common bioinformatics tools including fastp for sequencing read quality control and filtering, bwa for read alignment, samtools for option amplicon primer soft clipping and per-site depth calculations, and lofreq for assignment of indel qualities and variant calling [16–19]. Many of these pre-processing steps are wrapped into the integrated end-to-end workflow for convenience using snakemake [20]. External bioinformatics-focused Python libraries essential for the functioning of the tool include cyvcf2 for VCF file input, parsing and output, biopython for reference genome preparation, pandas and numpy for tabular processing and numerical work, matplotlib and seaborn for plotting, and the snakemake API for workflow orchestration [20–25]. Functional impacts of SARS-CoV-2 mutations are automatically inferred by searching the pokay database, which is a curated compendium of SARS-CoV-2 functional annotations and their connection to mutations [5].

### Inputs and required metadata

The main required input for vartracker is a CSV-format sample manifest detailing the input files, with the data type varying per analysis mode as described in the entry points description above. The manifest rows need to be ordered sequentially from the earliest to latest samples. A helper function to easily generate the input spreadsheet is executed via ‘vartracker prep spreadsheet [options]’.

Given its origins as part of a study investigating the evolution of SARS-CoV-2 throughout long-term serial passaging [3], vartracker comes packaged with SARS-CoV-2 assets. Therefore, if analysing data from SARS-CoV-2, vartracker can be deployed with no further amendments to the default parameters by relying on inbuilt SARS-CoV-2 reference assets and sensible analytical defaults. However, vartracker is generalisable to other pathogens if users provide suitable reference genome sequence (fasta format) and annotation data (GFF3 format). Compatible reference data can be generated from a GenBank accession string (or a file with accession numbers) using the inbuilt helper subcommand ‘vartracker prep reference’. Literature on the functional impacts of SARS-CoV-2 variants from the pokay database can be automatically downloaded if the ‘--search-pokay’ flag is provided, but for non-SARS-CoV-2 pathogens a compatible variant lookup table (CSV format) must be created and specified by the user. The expected database structure can be viewed using the ‘vartracker schema literature’ subcommand, and detailed instructions for creating the variant lookup table are provided in the project’s README (see ‘Building a custom literature database for other pathogens’)”.

### Processing and annotation workflow

The same downstream workflow is applied by vartracker across all analysis modes (Figure 1). Briefly, VCF files are first harmonised/normalised relative to the same reference genome and gene annotation to ensure equivalent mutations are described the same way in every sample. The amino acid consequences of mutations are then inferred using bcftools csq [16,26]. Per-sample results are then combined into one longitudinal table keyed by genomic position and allele, while retaining sample-specific measurements (allele frequency, variant-support depth, site/window depth, and QC flags). Variant-level quality control is then applied using configurable depth and allele-frequency thresholds (including separate SNV/indel cutoffs). A given variant is flagged as failing within a sample when no variant-supporting reads are present and site depth falls below the depth threshold, with mean depth across a 9 bp window either side of the affected span considered for indels. These QC parameters are incorporated to flag cases when genuine absence of a variant cannot be distinguished from dropout or non-detection. Flags are reported per sample, summarised per variant (all_samples_pass_qc, proportion_samples_passing_qc), marked on the default heatmap, and available as filtering criteria. yielding both per-sample QC and an overall QC status per variant. vartracker then applies longitudinal logic to classify variants by origin and persistence over time (baseline vs newly emerged; retained, lost, persistent, or transient), and records first/last appearance and presence/absence patterns across samples. Variants are also searched against a literature database if one is provided. Finally, outputs are summarised as tables and plots.

### Outputs and reproducibility features

The main longitudinal summarisation of variant data across samples is output as a schema-documented results table (CSV format). This table allows one to investigate easily both the biological insights and technical quality control from a longitudinal study, and filter quickly for high-value subsets of variants, such as newly arising variants that go to fixation over the course of the study. Additionally, vartracker flags if any of the called variants have hits within a provided literature database, outputting the hits within CSV-format output files and flagging variants within the output interactive HTML heatmap (see below).

Output plots generated by vartracker are straightforward to interpret and, with appropriate parameter choices, can be suitable for publication. Longitudinal variant allele-frequency heatmaps are produced as both static PDF files and interactive HTML documents, with optional links to published literature for detected mutations. Variant distributions across the genome are summarised in a PDF plot showing the genomic positions of variants together with their minimum and maximum allele frequencies. Gene-level mutation summary plots are also generated, showing total variant counts and counts by amino acid consequence class (for example, synonymous and missense), both with and without scaling by gene length. In addition, integrated plotting subcommands allow visualisation of variant trajectories, turnover, and lifespan over time based on patterns of presence or absence and changes in allele frequency. While several plots are generated automatically during the default workflow, they can also be regenerated and customised from an existing results file using ‘vartracker plot’ subcommands, allowing flexible replotting with customised parameters.

Each vartracker run records in a provenance file (JSON format) what was executed, with which software versions, and against which inputs/reference files. These records give a complete audit trail for reproducibility, making it easier to diagnose differences between runs. This supports transparent reporting in manuscripts and regulated workflows. Results are tied to an explicit schema version so downstream code can detect structural changes safely over time. The command-line interface schema command provides a machine- and human-readable description of output fields from the vartracker workflow, reducing ambiguity, improving interoperability, and making pipelines more robust to upgrades.

### Software availability, packaging and testing

vartracker is available as open-source code on GitHub (https://github.com/charlesfoster/vartracker), distributed under the permissive MIT license, enabling reuse, modification, and redistribution with minimal restrictions. The software is designed to be easily installed, most simply through its Bioconda distribution using conda, mamba, or pixi, or through corresponding container images provided via the Biocontainers registry [27]. vartracker requires Python ≥3.11 and is supported on Linux and macOS; pinned versions of all external tool dependencies are listed in the GitHub repository. Installable distributions are also packaged on PyPI, with a recommended conda environment to handle dependencies provided on GitHub. We also provide an optional Dockerfile to enable self-building of the package as a container. To support software quality and reproducibility, vartracker includes automated unit and command-line tests, and these are run through GitHub Actions workflows together with repository checks during continuous integration. Versioned releases provide stable, citable snapshots and clear change tracking across updates. Each formal release is archived with a Zenodo DOI, providing a permanent, version-specific record that supports reproducible analyses and precise citation of the software version used.

### Evaluation using representative datasets

Here, we demonstrate the practical utility of vartracker by analysing subsets of two published sequencing datasets. First, we analysed 13 samples from a long-term serial passaging experiment of a SARS-CoV-2 A.2.2 variant, representing whole-genome sequencing of isolates spanning the original clinical sample and 33 subsequent *in vitro* passages, with the aim of replicating a subset of the findings from the original study [3,28]. Raw reads were analysed using the end-to-end mode of vartracker with default settings, together with the ‘--search-pokay’ flag to enable automatic download and reformatting of the functional SARS-CoV-2 database from the pokay repository.

Second, to demonstrate that vartracker can also be applied to non-SARS-CoV-2 pathogens, we analysed sequencing data from five RSV-A samples from a longitudinal study investigating within-host human respiratory syncytial virus populations [29]. Reference files from the hRSV/A/England/397/2017 (PP109421.1) RSV-A reference genome were automatically downloaded and prepared using the ‘vartracker prep reference’ subcommand, after which the dataset was analysed in end-to-end mode using default settings apart from specifying the relevant reference inputs. Selected downstream plots were then regenerated using ‘vartracker plot’ subcommands to highlight features of interest, including the intersection of called variants with known antigenic sites within the F gene of RSV.

To assess the ability of vartracker to process data from a genome several orders of magnitude larger than a typical virus genome, we generated simulated longitudinal variant data for *Pseudomonas aeruginosa* PAO1. The PAO1 reference sequence and annotation (NC_002516.2; 6.26 Mb; 5,572 genes) were downloaded using ‘vartracker prep reference’. Coverage information and VCF files were simulated for six longitudinal timepoints under two scenarios: (i) a lower-variant-burden dataset containing 80 variants and (ii) a higher-variant-burden dataset containing 1000 variants concentrated in a small number of genes. The simulated inputs were processed using ‘vartracker vcf’ with default parameters besides the specification of the relevant reference genome files. Output accuracy was assessed by comparing the recovered variants and their presence or absence at each timepoint against the known simulated inputs. Annotation outputs and plots were also manually inspected to assess their interpretability.

### Computational environment and software provenance

All analyses were conducted with vartracker v2.3.0 on a MacBook Pro equipped with an Apple M5 Max chip, 18 CPU cores, and 64 GB unified memory, running macOS 26.4.1 on arm64 architecture. Exact commands for analysis of the SARS-CoV-2 and RSV-A datasets are provided in Supplementary File S1, and scripts and input data required to reproduce the *P. aeruginosa* simulations are available in the vartracker GitHub repository at scripts/validation/pao1.

Vartracker was conceived and implemented by the authors, and the source code developed through the initial stable release (v1.0.0) was written without generative-AI assistance. During subsequent development, the authors used GPT⍰5 and GPT⍰5⍰Codex (OpenAI) and Claude Opus 4.6 and Claude Sonnet 4.6 (Anthropic) to assist with codebase refactoring, minor bug fixes, packaging and distribution through PyPI, and scaffolding of new features. All AI-assisted code changes were reviewed by the authors through code inspection and validated using automated testing. The authors retained full responsibility for the design, implementation and validation of the software.

## Results and Discussion

### Performance on representative evaluation datasets

Across 13 samples in the SARS-CoV-2 serial passaging dataset, the mean per-sample depth of coverage was 1786.73× (median 1633.84×; range 1186.92×–3689.49×). The mean genome breadth of coverage at ≥10× was 99.61% (median 99.60%; range 99.55%–99.75%). End-to-end execution of ‘vartracker e2e’ on this dataset completed successfully in 4 min 29 s, with 1.69 GB peak RAM usage and an effective average CPU utilisation of approximately 3.3 cores across the full run. Our analysis using vartracker recapitulated key findings from the original study [3], including a putative partial haplotype replacement between passages 24 and 27, marked by the abrupt loss and gain of several near-fixed variants. Rapid shifts in the allele frequencies of key Spike gene variants linked to increased SARS-CoV-2 transmissibility and located around the furin cleavage site are shown in Figure 2, generated with the ‘vartracker plot trajectory’ subcommand.

**Figure 2.**
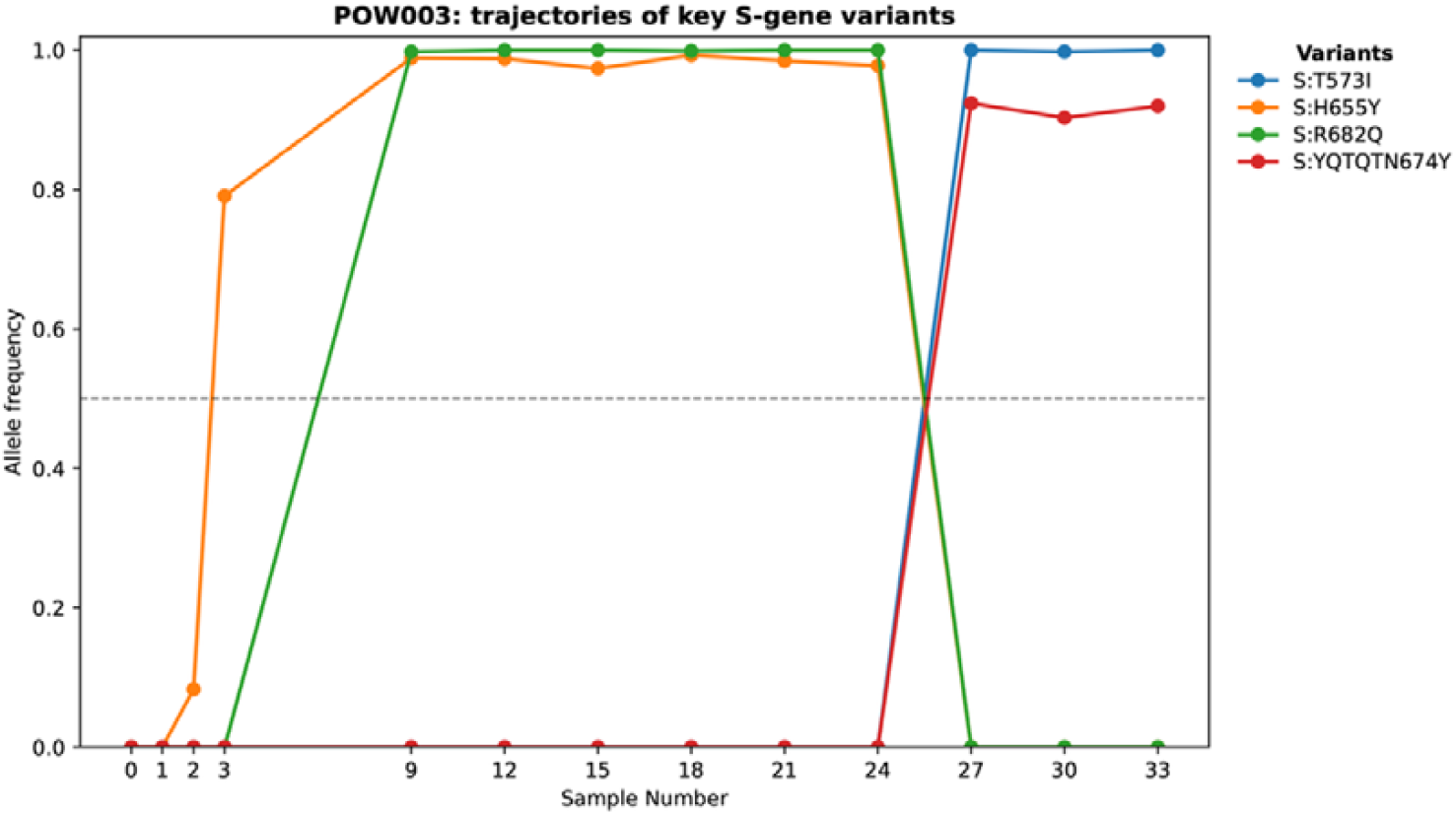
Longitudinal trajectories of key spike variants in the POW003 SARS-CoV-2 passaging series. Allele frequencies of selected spike variants are shown across serial passages from the original clinical sample (sample 0) to passage 33. The plot highlights a marked shift in the dominant variant profile between passages 24 and 27, with S:H655Y and S:R682Q declining and S:T573I and S:YQTQTN674Y emerging at high frequency. The dashed horizontal line indicates an allele frequency of 0.5. This plot was directly generated using the ‘vartracker plot trajectory’ subcommand, using data from [3].

The chosen RSV-A dataset was characteristically different from the SARS-CoV-2 dataset, with fewer samples (n = 5) at a much greater mean per-sample depth of 6639.03× (range 2863.02×–10490.45×). Genome breadth was 100% at ≥10× coverage in all samples, indicating complete callable coverage across the reference genome (PP109421.1). End-to-end execution on the RSV-A evaluation dataset (five samples) completed in approximately 7 min with 1.81 GB peak RAM usage and an effective average CPU utilisation of approximately 3.6 cores across the full run. Successful analysis of this dataset with vartracker provides proof of concept for its utility beyond SARS-CoV-2. The results recapitulated key findings from the original study [29], including detection of the minor variant G6406A, which corresponds to the inferred F:S255N amino acid substitution.

This substitution intersects with antigenic site II of the RSV F protein, which is the target for palivizumab and motavizumab antivirals. Using the integrated ‘vartracker plot genome’ subcommand, we plotted all detected variants across the F gene and specifically flagged those intersecting known antigenic sites (Figure 3), thereby partially replicating and extending Figure 1 of the original study [29].

**Figure 3.**
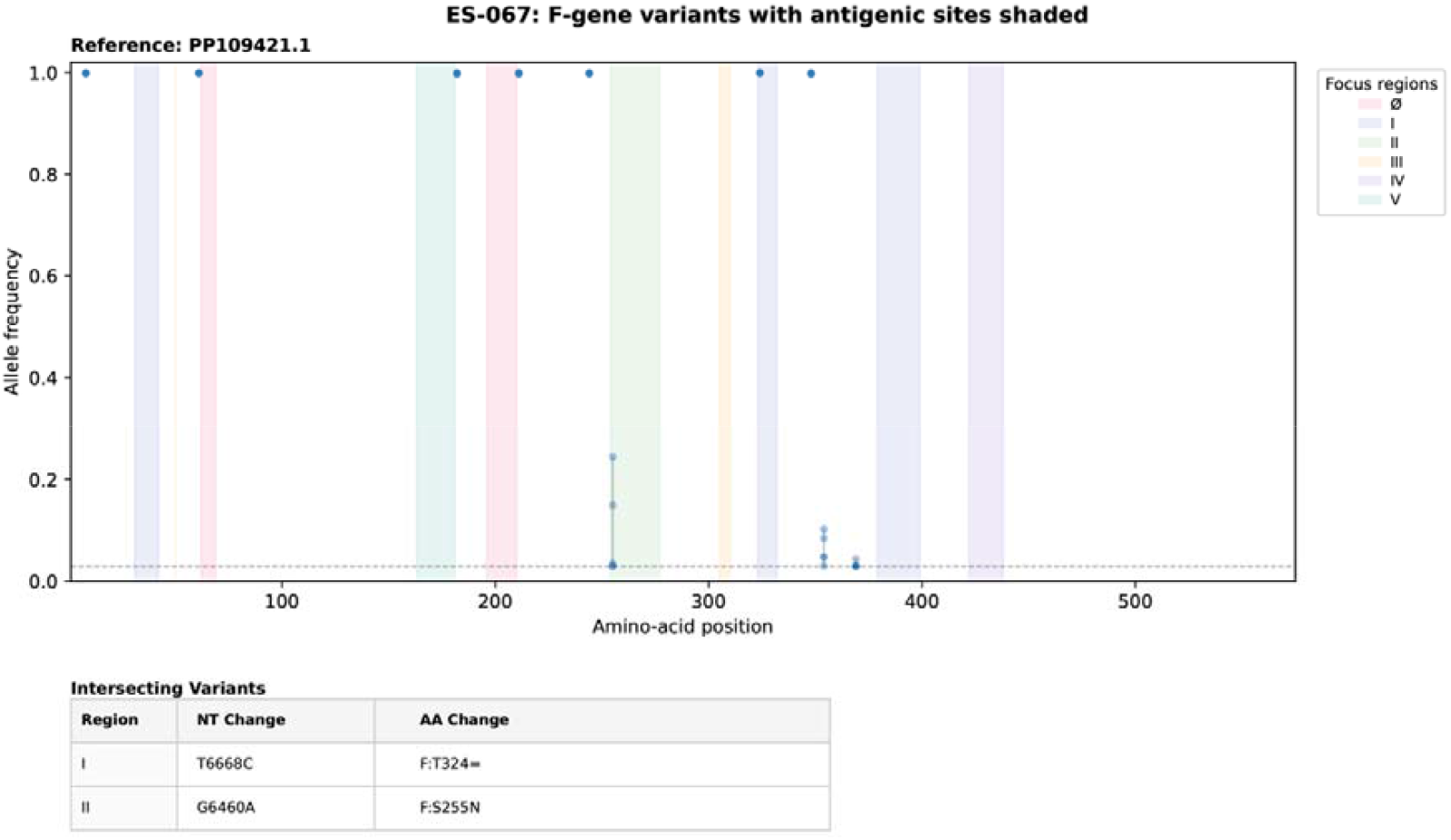
Distribution of F gene variants across a series of longitudinal RSV-A samples with antigenic sites highlighted. Variants identified in the F gene are plotted by coding-sequence position and allele frequency across the longitudinal ES-067 samples of [29]. Shaded vertical regions indicate selected antigenic sites in F, and the table summarises variants intersecting these highlighted regions. The dashed horizontal line marks the minimum allele-frequency threshold used for variant reporting. This plot was directly generated using the ‘vartracker plot genome’ subcommand.

Analysis of a simulated bacterial dataset containing 80 variants across six timepoints using ‘vartracker vcf’ completed in approximately 11 seconds, with peak memory usage of approximately 0.9 GB. All simulated variants and their expected temporal patterns were recovered, and the resulting plots were readily interpretable. Analysis of a higher-complexity dataset containing 1000 variants across six timepoints completed in approximately 22 seconds, with peak memory usage of approximately 2.8 GB. All expected variants and their temporal presence/absence patterns were recovered correctly at the nucleotide level, but the “joint” amino-acid consequences reported by ‘bcftools csq’ were harder to interpret than in the 80-variant simulation or compared to a typical viral analysis.

Vartracker runs ‘bcftools csq’ in non-reference-haplotype mode (’--phase R’) because its genotypes are unphased by construction, which causes co-occurring variants within a sample to be treated as though linked on the same haplotype. Because the specific set of co-occurring variants can then differ from timepoint to timepoint, a single variant can accumulate several distinct, lengthy compound consequence descriptions across the series. However, this annotation complexity is a readability issue at the amino acid level rather than a correctness one: nucleotide-level tracking of variant presence or absence is unaffected since it is independent of the consequence descriptions. To improve readability, users can instead invoke vartracker with the ‘--local-csq’ flag, which predicts consequences for individual variants independently and produces shorter annotations similar to those generated by SnpEff or equivalent tools. The trade-off is that genuine combined effects between physically linked variants are not captured. For example, two indels that would individually be predicted to cause frameshifts may jointly restore the reading frame, but independent annotation would not represent this combined effect. Users can also filter the variants displayed by the ‘vartracker plot’ subcommands when visualising datasets containing many variants.

### Practical utility for annotation and interpretation

vartracker simplified analysis and interpretation in the experimental comparator sets, from raw reads to downstream gene-and amino-acid-aware summaries and visualisations, within a single command. In the SARS-CoV-2 dataset, the ‘vartracker plot trajectory’ subcommand enabled rapid inspection of coordinated allele-frequency shifts across passages, a pattern that might otherwise have been less readily apparent. Optional literature-aware annotation further assisted prioritisation of potentially important variants, while the resulting interactive heatmap visualised their allele-frequency trajectories and provided direct links to supporting literature. For the RSV-A dataset, preparation of suitable reference files was streamlined using the ‘vartracker prep reference’ subcommand. This step was particularly useful because it generated reference annotations compatible with bcftools csq for amino acid consequence inference, whereas default GFF3 files from NCBI GenBank are often not fully compatible. The ‘vartracker plot genome’ subcommand also highlighted variants intersecting predefined antigenic regions and generated a publication-ready plot. Together, these examples illustrate the practical value of vartracker as a lightweight interpretation layer for longitudinal pathogen variant data.

### Intended use and limitations

The primary intended use for vartracker is to efficiently summarise, annotate and visualise longitudinal variant calling results. vartracker is therefore designed as a data-exploration tool for interrogating the biological implications of longitudinal studies, rather than as a primary variant-calling method. Its outputs consequently depend on the quality of upstream alignment, consensus generation and variant-calling steps, and on the consistency of that processing across the samples in a series. Where serial samples derive from different library preparations, platforms, basecallers or amplicon schemes, the resulting differences in coverage profile are captured by the per-sample quality control flags described above. However, differences in caller sensitivity and allele-frequency estimation at adequately covered sites are not, since vartracker applies uniform thresholds across a series and does not model or correct for between-source technical variation. Origin and persistence classifications therefore describe the supplied call set, and users pooling data from heterogeneous sources should process samples through a single consistently configured workflow where possible, and treat emergence or loss events that coincide with a change in source as provisional.

The e2e and bam entry modes of vartracker are intended as convenient integrated workflows rather than definitive variant-calling solutions. Performance in these entry modes will vary with dataset characteristics such as sample number and sequencing depth. In particular, very deeply sequenced datasets may be bottlenecked by the integrated variant-calling step, as was observed when analysing the more deeply sequenced RSV-A dataset in end-to-end mode. While the e2e and bam modes offer reasonable flexibility, users with specific upstream processing or variant-calling preferences may be better served by calling variants separately and supplying these to vartracker in vcf mode. This separation is intentional, as the main novelty of vartracker lies not in replacing established variant-calling methods, but in the downstream interpretation layer it provides. Specifically, vartracker adds value through its longitudinal summarisation of variants across serial samples, integrated amino-acid and literature-aware annotation, and purpose-built plotting functions for exploring temporal trajectories and genomic context. In this way, the tool is designed to complement, rather than compete with, upstream variant-calling workflows.

Vartracker is intended primarily for longitudinal studies involving several to tens of ordered samples from an individual, experimental system or defined outbreak investigation, rather than population-scale surveillance involving thousands of samples. To gauge the scalability of vartracker, in addition to the case studies presented above, we also analysed longitudinal passages 0 to 100 of the POW005 SARS-CoV-2 passage line (37 samples total) from Foster et al. [3,28]. The end-to-end workflow successfully processed this dataset in approximately 9 min 43 s using 16 allocated CPU cores with peak single-process memory usage of 1.7 GB, whereas the lightweight vcf entrypoint, reusing the resulting VCF and coverage files, completed in only 15 s with peak single-process memory usage of 1.2 GB (excluding download of the pokay database, which added approximately 34 s). Although these analyses do not establish a computational upper limit for vartracker, they demonstrate practical use beyond the 5–13-sample series examined in the principal case studies. Performance on substantially larger cohorts has not been systematically evaluated. A practical further limitation for users to consider is the legibility of generated plots with increasing numbers of samples. In these circumstances, users are encouraged to explore the filtering and customisation parameters of the ‘vartracker plot’ subcommands to optimally generate figures.

Further evaluation across larger and more heterogeneous longitudinal datasets, together with systematic comparisons against existing analysis and visualisation platforms, will be valuable for characterising the scalability and utility of vartracker across broader applications. The principal value of vartracker lies in enabling biologically meaningful exploration of longitudinal viral variation across outbreak investigations, experimental studies and within-host analyses, where temporal patterns of mutation may have evolutionary, immunological, clinical or public health relevance.

## Data Availability

The source code for vartracker is released under the MIT License and is available from the GitHub repository charlesfoster/vartracker at https://github.com/charlesfoster/vartracker. Versioned releases are archived on Zenodo at https://doi.org/10.5281/zenodo.18452274. The raw sequencing data for SARS-CoV-2 and RSV-A analysed in this study were obtained from the NCBI Sequence Read Archive (BioProject PRJNA1018257) and the European Nucleotide Archive (Project PRJEB34042), respectively.

